# Enhanced recovery of single-cell RNA-sequencing reads for missing gene expression data

**DOI:** 10.1101/2022.04.26.489449

**Authors:** Allan-Hermann Pool, Helen Poldsam, Sisi Chen, Matt Thomson, Yuki Oka

## Abstract

Droplet-based 3’ single-cell RNA-sequencing (scRNA-seq) methods have proved transformational in characterizing cellular diversity and generating valuable hypotheses throughout biology^1,2^. Here we outline a common problem with 3’ scRNA-seq datasets where genes that have been documented to be expressed with other methods, are either completely missing or are dramatically under-represented thereby compromising the discovery of cell types, states, and genetic mechanisms. We show that this problem stems from three main sources of sequencing read loss: (1) reads mapping immediately 3’ to known gene boundaries due to poor 3’ UTR annotation; (2) intronic reads stemming from unannotated exons or pre-mRNA; (3) discarded reads due to gene overlaps^3^. Each of these issues impacts the detection of thousands of genes even in well-characterized mouse and human genomes rendering downstream analysis either partially or fully blind to their expression. We outline a simple three-step solution to recover the missing gene expression data that entails compiling a hybrid pre-mRNA reference to retrieve intronic reads^4^, resolving gene collision derived read loss through removal of readthrough and premature start transcripts, and redefining 3’ gene boundaries to capture false intergenic reads. We demonstrate with mouse brain and human peripheral blood datasets that this approach dramatically increases the amount of sequencing data included in downstream analysis revealing 20 - 50% more genes per cell and incorporates 15-20% more sequencing reads than with standard solutions^5^. These improvements reveal previously missing biologically relevant cell types, states, and marker genes in the mouse brain and human blood profiling data. Finally, we provide scRNA-seq optimized transcriptomic references for human and mouse data as well as simple algorithmic implementation of these solutions that can be deployed to both thoroughly as well as poorly annotated genomes. Our results demonstrate that optimizing the sequencing read mapping step can significantly improve the analysis resolution as well as biological insight from scRNA-seq. Moreover, this approach warrants a fresh look at preceding analyses of this popular and scalable cellular profiling technology.

## Main

Droplet-based single-cell RNA-sequencing methods such as Dropseq and 10x Genomics platforms have dramatically lowered the cost and improved the throughput of single-cell gene expression profiling. These advances have thereby widely democratized the discovery of new cell types and states^6–8^, delineation of developmental mechanisms^9^ and cellular basis of disease^10^ as well as mapping of behavioral and physiological functions to distinct cell types^11,12^. The scalability of such methods however comes with a few important limitations. First, the droplet-based methods rely on 3’ gene tagging where detection of genes depends on registering sequencing reads predominantly at the 3’ end of genes which makes detection of splicing isoforms problematic. Second, 3’ scRNA-seq datasets despite usually being much more shallowly sequenced are in general considered to have lower sensitivity than deep full-length isoform sequencing solutions such as provided by the SMART-Seq chemistry^13^. Indeed, several studies have observed that genes shown to be expressed with other methods have critically been missing in analyses relying on droplet-based scRNA-seq^8,11^. This shortcoming compromises the potential of 3’ scRNA-seq high-throughput technologies to uncover the genetic and cellular mechanisms giving rise to development and tissue function.

ScRNA-sequencing workflow consists of several steps including sample preparation, sequencing library generation, sequencing, read mapping/quantification, and analysis of the gene-cell matrix based data. While many of these steps are considered standard, some such as sample preparation are widely recognized as critical for the final outcome and can vary significantly between protocols and labs. One often overlooked step in this workflow is read mapping/quantification that determines which sequencing reads are incorporated in the final cellular gene expression data. During this process, sequencing reads are mapped to the reference transcriptome (i), assigned to genes (ii), assigned to cells (iii), and duplicates are removed (iv) ^14,15^. As a result of this step, often the majority of sequencing reads get excluded from further analysis for one of several reasons including failure to map confidently to the transcriptome, being a duplicate read, mapping to multiple sites in the genome (multimapping reads), mapping to more than one gene (multigene reads), mapping intronically or to an intergenic region. Some of the discarded read data however reflect endogenous gene expression and can render expressed genes missing^16,17^. Several groups have manually amended the transcriptome for individual genes to restore their visibility^8,11^, however, a systemic effort to evaluate the scale of this problem and to provide a whole-transcriptome solution for this issue has been missing.

Here, we show that analysis pipelines relying on standard exonic transcriptomic references are blind to many genes that are easily detected with independent methods such as in situ hybridization. We demonstrate that this lack of gene detection does not stem from low sensitivity but rather inefficiencies of the currently used transcriptomic references and that this is the case even with very well annotated genomes including that of mouse and human. Furthermore, we show that the read loss stems from three sources: poor annotation of 3’ untranslated regions, gene overlaps stemming from the annotation of rare read-through or prematurely starting transcripts and finally exclusion of intronic reads. We outline a three-step strategy to overcome these limitations through the inclusion of intronic reads, resolving gene overlaps by excluding rare transcript isoforms and identifying and incorporating unannotated gene 3’UTRs. This strategy recovers obscured gene expression data for thousands of genes and reveals previously undetected genetic markers, mechanisms and cell types. Consequently, we provide full genome optimized transcriptomic references for the mouse and human genomes. In sum, our data argue that transcriptomic references need to be optimized for scRNA-seq analysis and that this step can dramatically improve the profiling resolution. These findings also warrant a reanalysis of previously published datasets.

## Results

In order to characterize gene detection fidelity of 3’ gene counting methods we performed scRNA-sequencing of the median pre-optic nucleus (MnPO) - a mouse brain center implicated in a range of physiological functions including thirst, sleep, heat and cold sensation^18^. Predictably, following sequencing read mapping to an exonic transcriptomic reference we identified about a dozen distinct neuron types in this structure reflecting the functional diversity of this brain center (Fig. 1a). We next compared gene detection fidelity with scRNA-sequencing to in situ hybridization – an independent method provided by the Allen in situ brain atlas^19^. While we found many genes that were reliably detected with both methods (e.g. Nxph4, Fig. 1b), we observed a number of genes that were completely missing in scRNA-seq data while robustly detected with in situ hybridization (e.g. B4galnt2 and Gpr165, Fig. 1c-d). Follow-up analysis at these loci revealed three distinct patterns of sequencing read mapping that determined whether the gene is detected or missing in scRNA-sequencing analysis. The first type comprised of genes detected by both methods. In this case, sequencing reads mapped near perfectly to the exons of the underlying gene and were thus included in downstream transcriptomic analysis (Fig. 1b). A second group of genes were detected by in situ hybridization but were missing in scRNA-seq data as most sequencing reads mapped to an intron of that gene resulting in exclusion from transcriptomic analysis (Fig. 1c). Finally, a third group of genes were detected by in situ hybridization but not with scRNA-seq and had no sequencing read mapping to known exons and introns (Fig. 1d). Importantly, the last type of genes displayed excessive read mapping proximal to the known 3’ end of the gene suggesting that scRNAseq fails to detect these genes due to poor annotation of 3’ untranslated regions of genes. These data demonstrate that droplet-based single cell sequencing datasets can fail to detect genes due to suboptimal read mapping to the reference transcriptome.

**Figure 1:**
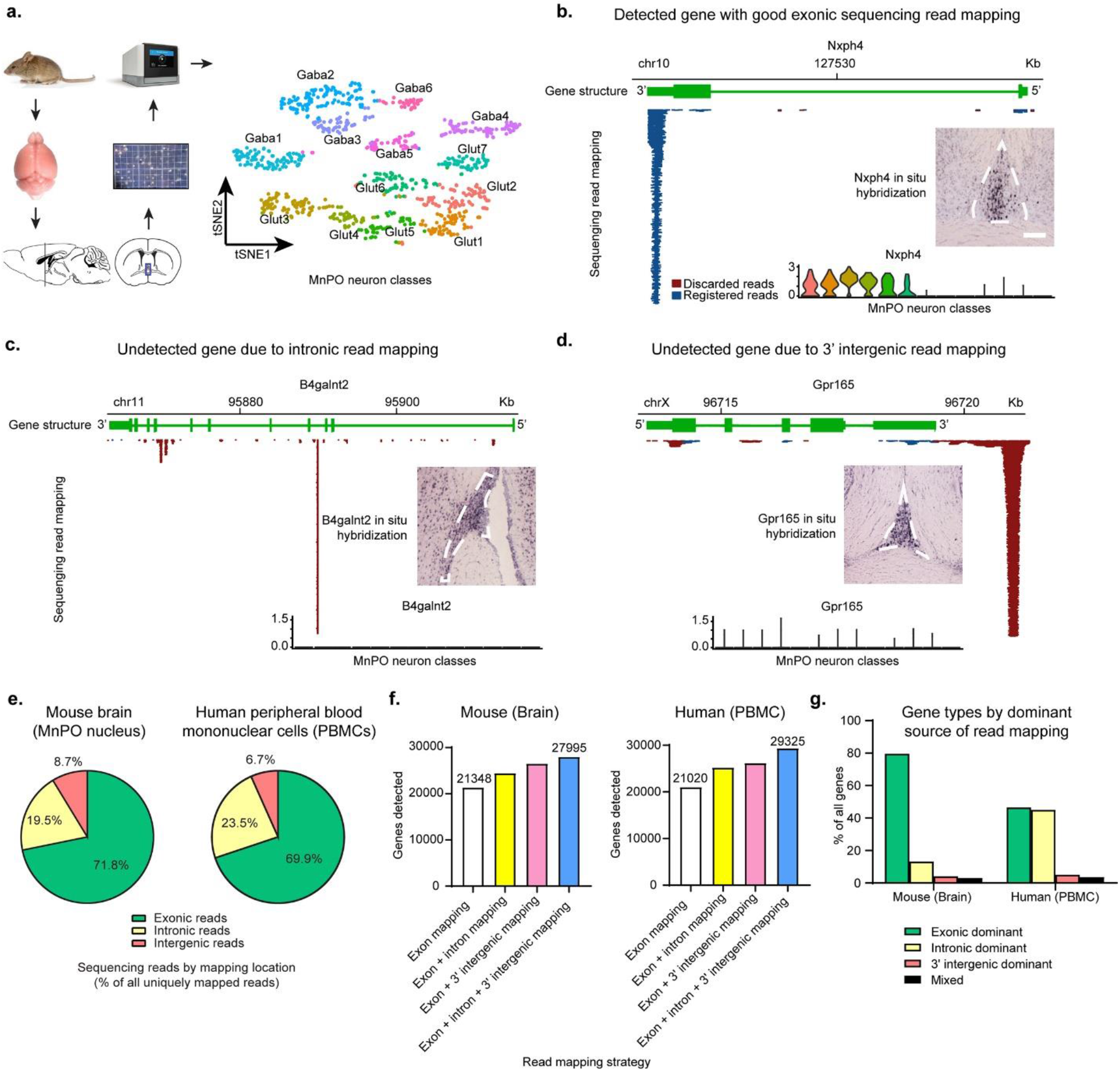
Missing genes and sequencing read registration in single-cell RNA-seq experiments. **a.** Sc-RNA-seq based profiling of the mouse physiology regulating brain center - Median Preoptic Nucleus (MnPO). 10x Genomics 3’ transcriptomic analysis of MnPO neurons (n=906) mapped to an exonic transcriptomic reference reveals 13 neuron types. Data shown in a tSNE embedding. **b.** Sample scRNA-seq detected gene (Nxph4) with sequencing read mapping at its genomic locus. The majority of sequencing reads map to known exons of Nxph4 gene and are therefore registered (blue) and included in downstream analysis. Discarded reads (red) map to non-exonic regions or are antisense to the gene and are therefore excluded. Inset violin plot: scRNA-seq analysis detects Nxph4 expression in several MnPO neuron types (cell-type specific log-transformed expression of Nxph4 in MnPO neuron types with cell-type identity color-coded as in Fig1a). Micrograph inset: in situ hybridization of Nxph4 expression in the MnPO (scale bar: 150 µm, posterior MnPO outlined with white dashed line, data from Allen Brain Atlas Mouse ISH dataset). **c.** Sample gene (B4galnt2) not detected by scRNA-seq due to intronic read mapping. Inset violin plot: gene expression is not detected in any of the MnPO neuron types. Inset micrograph: in situ hybridization of B4galnt2 expression in the MnPO. **d.** Sample gene (Gpr165) not detected by scRNA-seq due to intergenic read mapping 3’ of known end of the gene. Inset violin plot: gene expression is not detected in any of the MnPO neuron types with scRNA-seq. Inset micrograph: in situ hybridization of Gpr165 expression in the MnPO. **e.** Proportion of uniquely mapped sequencing reads according to mapping site (exonic, intronic or intergenic) for mouse brain (MnPO, left) and human peripheral blood mononuclear cells (right) datasets. **f.** Intronic and intergenic reads constitute a promising source to recover missing gene expression data in scRNA-seq analysis. Number of detected genes in mouse brain (MnPO, left) and human PBMC (right) datasets, if reads mapping to exons, exons and introns, exons and intergenic reads within 10kb of known 3’ ends of genes, or all three sources are included in downstream analysis. **g.** Human and mouse genes according to the dominant source of sequencing read data. Genes are classified as ‘exonic dominant’, ‘intronic dominant’ or ‘3’ intergenic dominant’ if more than 50% of sequencing reads map to their exons, introns or within 10kb of their 3’ end, respectively. Mixed genes have less than 50% of reads stemming from any of the three regions.

In order to evaluate the magnitude of the missing gene problem, we quantified several metrics of sequencing read mapping in two vertebrate species with the most thoroughly annotated genomes – mice and humans. For mice we evaluated the MnPO dataset and for humans we profiled peripheral blood mononuclear cells (PBMCs). In mouse brain data we found that out of the uniquely mapped sequencing reads 71.8 % are exonic, 19.5 % intronic and 8.7 % intergenic out of 272 million total reads suggesting that significant gains could be achieved by incorporating sequencing data from intronic and intergenic areas to gene expression estimates (Fig. 1e). We found similar metrics in human data with 69.9 % exonic, 23.5 % intronic and 6.7 % intergenic reads (272 million total), respectively. Indeed, upon evaluating the number of genes detected as a result of including intronic reads, intergenic reads within 10 kb of known 3’ gene ends or both, we observed dramatic gains in the amount of detected genes in scRNA-seq datasets with 13.6%, 25.8% and 33.6% more genes detected than with a conventional exonic transcriptome reference in mouse (Fig. 1f). Again, comparable gains were observed with 19.9%, 23.2% and 39.2% more genes detected, respectively for the human transcriptome. Moreover, we also evaluated the dominant source of read information for genes in the mouse and human datasets. Predictably we found that the majority of mouse genes (79.6%) were dominated by exonic reads with more than 50% of expression data stemming from exonic reads (Fig. 1g). Somewhat surprisingly, less than half of human genes derive their expression data from exonic reads with the rest stemming from intronic or 3’ intergenic reads. While not all intronic and proximal intergenic sequencing reads stem from the respective protein-coding gene transcripts, these data indicate that profound gains in gene detection sensitivity are feasible by incorporating relevant intronic and intergenic read data in downstream scRNA-seq analysis.

We further evaluated the extent to which intergenic reads 3’ from gene ends could contribute to true gene expression estimates. If unannotated 3’ UTRs constitute a significant source of read loss in 3’ scRNA-seq datasets we would expect to see elevated levels of sequencing reads mapping proximal to 3’ end of genes. Indeed, we observe several-fold higher mapping of intergenic reads immediately proximal to the 3’ gene ends than at distal sites in both mouse and human datasets (Fig. 2a, b). In fact close to 25% of intergenic reads in both mouse and human datasets are within 10kb of 3’ gene ends, which represents approximately two-fold enrichment as compared to the rest of the non-coding genome^20,21^. These results suggest that improved annotation of 3’ gene ends is a promising strategy to increase gene detection in 3’ single-cell RNA-sequencing analysis (Fig. 2c).

**Figure 2:**
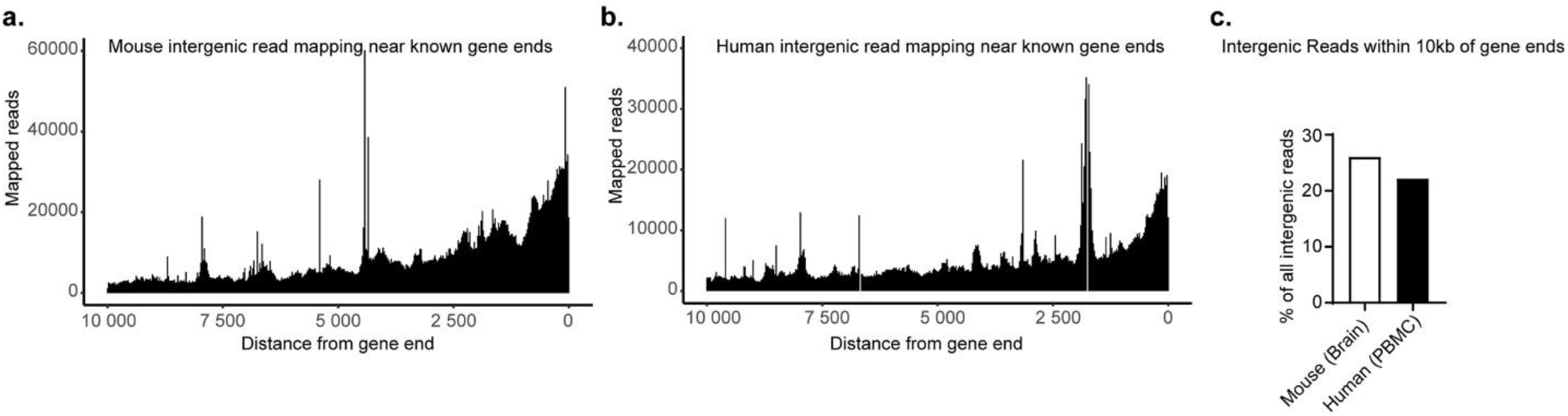
Increased intergenic read mapping proximal to 3’ end of genes. **a.** Distribution of sequencing reads mapping within 10kb of known gene ends in the mouse genome shows increased mapping proximal to gene ends. **b.** Distribution of sequencing reads mapping within 10kb of known gene ends in the human genome shows increased mapping proximal to gene ends. **c.** Fraction of intergenic reads mapping within 10kb of known gene ends from all intergenic reads in the mouse brain (MnPO) and human PBMC datasets.

Another common source of read loss in scRNA-seq analysis stem from same strand gene overlaps. Reads mapping to genomic regions annotated to more than one gene are classified as multigene reads and are routinely removed from downstream analysis^14,15^. We evaluated the magnitude of gene overlaps using the Ensembl mouse (v.98) and human (v.98) genome annotations which are most commonly used to generate reference transcriptomes for scRNA-seq analysis. We found that gene overlaps are a pervasive feature of currently available genome annotations with 2035 (6.3 % of all mouse genes) and 5195 (14.2% of all human genes) genes showing partial or complete overlap with other same strand genes in the mouse and human genomes, respectively (Fig. 3a). The majority of these overlaps in both mouse and human genomes originate from single pairs of genes (Fig. 3b, c).

**Figure 3:**
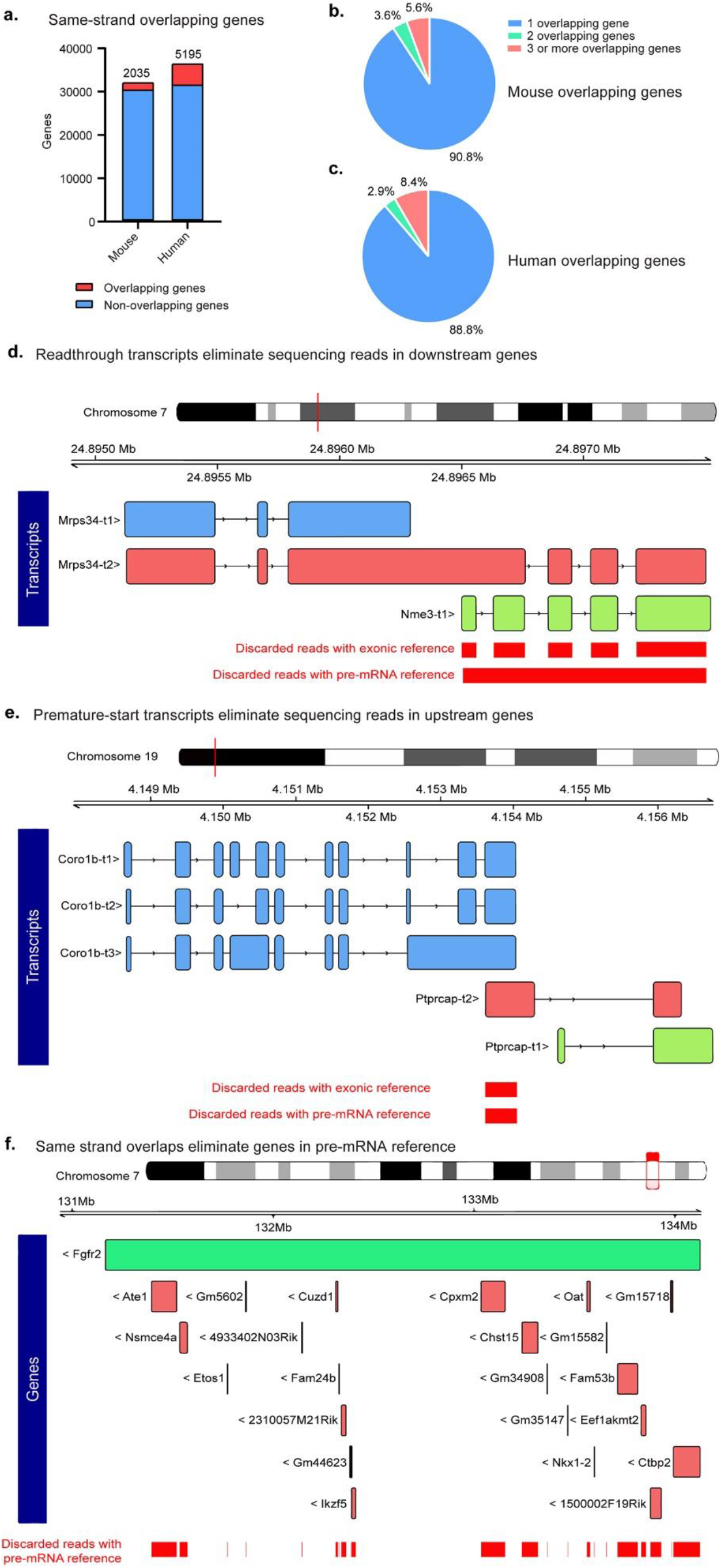
Gene overlap and resulting compromised scRNA-seq gene detection in the mouse and human genomes. **a.** Number of same-strand overlapping genes in the mouse and human genomes (mouse annotation - Ensembl v98 for GRCm38 build; human annotation - Ensembl v98 for GRCh38 build). **b.** Number of gene overlaps among mouse overlapping genes. **c.** Number of gene overlaps among human overlapping genes. **d.** Readthrough transcripts prevent the incorporation of sequencing reads to gene expression estimates in downstream genes. Gene regions where sequencing read data are discarded from gene expression estimates due to multigene classification are highlighted in red. **e.** Premature-start transcripts prevent the incorporation of sequencing reads to upstream gene’s expression estimates. Gene regions where sequencing read data are discarded due to multigene classification are highlighted in red. As most sequencing reads map at the 3’ end of genes, premature-start transcripts can render upstream genes undetectable by scRNA-seq analysis. **f.** Large multiple gene spanning genes can eliminate scRNA-seq detection of dozens of nesting same-strand overlapping genes depending on read mapping strategy. With pre-mRNA references, where full gene spans are defined as exons, all nesting genes will have no sequencing reads incorporated into expression estimates due to resulting multi-gene mapping classification.

A closer inspection of overlapping genes revealed a few stereotypic patterns of overlaps that result in partial or complete blinding of one or more overlapping genes from downstream analysis. The first problematic pattern stems from readthrough transcripts where one or several of upstream gene’s transcripts incorporate some or all exons of a downstream gene which effectively eliminates all sequencing reads mapping to the latter (Fig. 3d). Another problematic feature of overlapping genes are so called „premature start transcripts” where a single or several transcripts from a downstream gene are annotated to start upstream of the upstream gene’s terminal exon (Fig. 3e). The latter type of overlap is particularly problematic as the majority of sequencing reads in 3’ scRNA-seq map to terminal exons and thus premature start transcripts effectively eliminate the entire detection of their upstream gene. A version of this issue impacts dozens of genes that share their terminal exon and are thus completely invisible to analysis (Extended Fig. 1). Finally, multigene overlapping genes pose a particular problem for pre-mRNA references where a single large gene can completely eliminate dozens of nested genes rendering downstream analysis blind to their expression (Fig. 3f). An important caveat to the latter is that there are currently several strategies for compiling a pre-mRNA transcriptomic reference with substantial differences in gene detection and read mapping fidelity (Extended Fig. 2). In summary, gene overlaps in genome annotations constitute a unique challenge to discovering valuable candidate genetic mechanisms and marker genes in 3’ single-cell RNA-seq analysis. Moreover, these problems impact thousands of genes particularly in well annotated genomes.

The systemic issues with read loss stemming from discarding intronic, intergenic and multigene mapping reads outlined above (Fig. 4a) suggest a straight-forward strategy to optimize transcriptomic references. Here, we implement a three step process to overcome these limitations that is applicable for any genomic annotation. In the first step we convert an exonic reference to a pre-mRNA reference to incorporate intronic reads into gene expression estimates using a hybrid intronic mapping strategy (Fig. 4b). Secondly, we resolve gene overlaps by automated identification and curation of premature and readthrough transcripts eliminating overlapping transcripts, gene models and long non-coding RNA genes that obscure or preclude detection of protein-coding genes (Fig. 4c). Finally, we incorporate unannotated 3’ UTRs into our gene models by rank ordering genes with high sequencing read mapping within 10kb of their known gene end and supervised 3’ gene extension based on one of several criteria: a) read splicing to known exons, b) extended gene boundary in another genome annotation (e.g. Refseq), c) external ground truth evidence (Allen in situ atlas, Protein Atlas etc). As a result we generated optimized genome annotations for both mouse and human transcriptomes (Fig. 4e, Suppl. Tables 1, 2). This constitutes a general and scalable strategy for optimizing genome annotations for high-efficiency 3’ scRNA-seq analysis.

**Figure 4:**
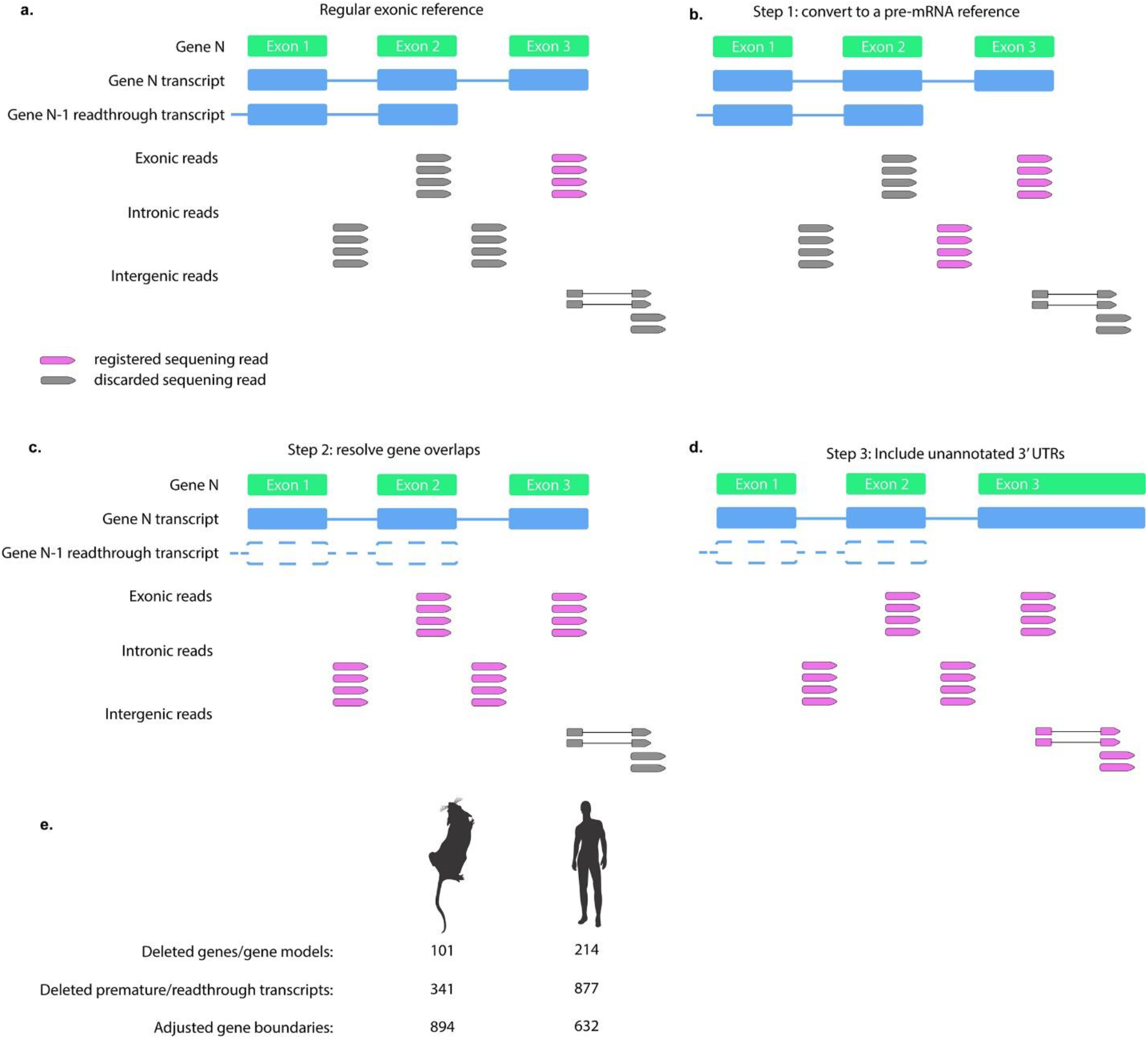
Strategy for compiling an optimized transcriptomic reference. **a.** Schematic of read registration with regular exonic reference. Registered sequencing reads that are incorporated to gene expression estimates are highlighted in purple with discarded sequencing reads shown in grey. ScRNA-seq analysis with an exonic reference discards several types of sequencing reads that map to a specific gene including intronically mapped reads, reads mapping to exons that overlap with readthrough transcripts from upstream genes (N-1) as well as sequencing reads mapping to unannotated 3’ untranslated regions (UTRs). **b.** Step 1 of optimizing a transcriptomic reference is incorporating intronic reads thereby generating a “pre-mRNA reference”. **c.** Step 2 of optimizing a transcriptomic reference is resolving gene overlaps by removing rare readthrough and premature transcripts as well as poorly supported gene models and pseudogenes that result in eliminating sequencing data from well-established protein-coding genes. This step incorporates sequencing reads mapping to exons and introns that previously overlapped with readthrough/premature transcripts. **d.** Step 3 of optimizing a transcriptomic reference entails extending 3’ boundaries of genes to incorporate unannotated 3’ UTRs with sequencing reads spliced to reads mapping to known exons. **e.** Genome annotation modifications for optimized mouse and human reference transcriptomes.

In order to evaluate the performance of the optimized reference transcriptomes, we evaluated the gene and read detection efficiencies in both mouse brain and human PBMC datasets, and contrasted the analyses to the same scRNA-seq dataset mapped to the traditional exonic reference. We observed dramatic gains in both gene detection and read registration with the optimized mouse transcriptome with more than 3000 new detected genes and 14.8% more sequencing reads included in downstream analysis. Moreover, the optimized reference yields a profound increase in cellular profiling resolution with close to 600 additional genes/cell on a median basis for MnPO neurons that constitutes a more than 20% increase in the number of genes detected per neuron (Fig. 5a). Furthermore, this increase in cellular profiling resolution translated into 1 – 3 additional neuron types detected under identical analysis parameters to exonic transcriptome based analysis. Predictably, the optimized transcriptome revealed genes that were invisible to the traditional exonic reference based scRNA-seq analysis due to sequencing read mapping to intronic and un-annotated 3’ UTR reigons (Fig. 5b).

**Figure 5:**
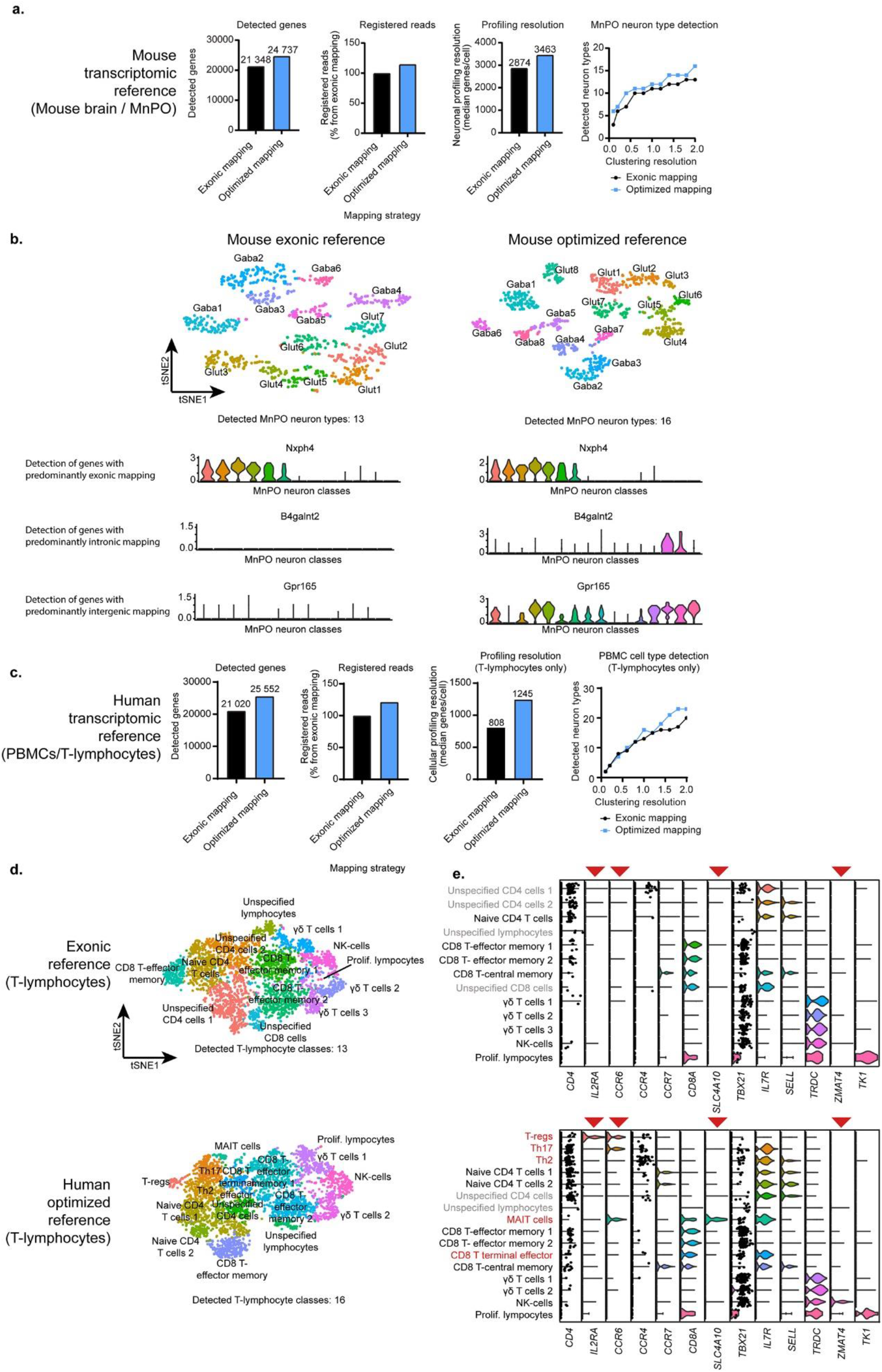
Evaluation of optimized mouse and human reference transcriptomes. **a.** Quantification of the mouse exonic and optimized transcriptomic reference performance with the MnPO scRNA-seq dataset. Detected genes and registered reads are estimated from the raw gene-cell matrix produced by Cell Ranger count pipeline using the canonical exonic reference and the optimized reference, respectively. Profiling resolution is estimated as median genes detected per cell based on the same set of neurons (n=906) in the MnPO dataset. Neuron type detection is estimated on the same set of neurons with identical preprocessing parameters with the x-axis denoting the clustering granularity hyper-parameter for the Louvain algorithm for modularity optimization. **b.** Comparison of scRNA-seq analysis with exonic and optimized transcriptomic references as deployed to MnPO neurons (n=906). Optimized reference reveals more neuron types at identical data preprocessing conditions. Optimized reference also detects sample genes with intronic (B4galnt2) and 3’ intergenic (Gpr165) read mapping that remain invisible with scRNA-seq analysis relying on the exonic transcriptomic reference. **c.** Quantification of the human exonic and optimized transcriptomic reference performance with the peripheral blood mononuclear cell (PBMC) data. Gene detection and read registration estimates stem from full PBMC data evaluation. Cellular profiling resolution and cell-type detection is estimated on the same set of T-lymphocytes extracted from the PBMC dataset (n= 4133) with identical preprocessing parameters. **d.** Transcriptomic analyses of the subtypes of T-lymphocytes (n=4133) in PBMC data mapped to the exonic reference (above) or optimized human reference transcripome (below). ScRNA-seq data are shown in a tSNE embedding of 4133 lymphocytes with color-coded cell identity. **e.** Violin plots of log-normalized expression data of T-lymphocyte subtype marker genes shown for PBMC data mapped to the exonic reference (above) or optimized human reference transcriptome (below). Ambiguous and cell classes newly detected with the optimized transcriptomic reference are color coded gray and red, respectively. Canonical cell class specific markers that are missing in exonic reference analysis and recovered in optimized reference analysis have been highlighted with red triangles.

We found consistently superior performance of the optimized human genome annotation based analysis as compared to the implementation of an exonic transcriptomic reference. We detected over 4500 additional genes and more than 21% of additional sequencing reads in the human PBMC dataset (Fig. 5c). Similarly to the optimized mouse transcriptome, we observed dramatic gains in profiling resolution of cells with more than 400 additional genes/cell detected on a median basis. These gains in gene and read detection in the human dataset translated to up to 6 additional cell types detected under identical analysis parameters as compared to the analysis based on the exonic trancriptomic reference. Therefore, optimizing genome annotations for scRNA-seq analysis can lead to robust gains in sequencing read, gene as well as cell-type detection.

The key value of implementing an optimized transcriptomic reference is to reveal biological hypotheses and pinpoint genetic markers that would otherwise remain unobserved. To test the merit of optimizing transcriptomic references we sought to compare and contrast the performance of exonic to the optimized transcriptomic reference in linking MnPO cell types to their underlying physiological functions. MnPO has been implicated in a range of physiological functions including thirst, heat stress, cold stress, sleep and licking^18^. While a comprehensive single cell profiling of MnPO has not previously been performed, a number of studies have attempted to find genetic markers labeling neurons mediating the previous functions. Thus, MnPO neurons expressing Vglut2, Nos1, Nxph4 and Pacap^22–25^ have been implicated in thirst regulation; neurons labeled by Pacap, Bdnf, Ptger3 and Trpm2 have been suggested to label MnPO warmth induced neurons ^26–28^; Glp1r+ neurons have been shown to be selectively driven by licking and liquid ingestion^23^; and finally Brs3+ neurons have been shown to label cold activated neurons in the MnPO^29^.

Neuron type resolved view of these markers in the MnPO (Extended Fig. 3a, b) revealed that all the thirst markers are widely expressed in half or more of the cell types suggesting that these are unlikely to be specific for this function. Moreover, we observed that while two of the putative heat stress markers (Bdnf and Pacap) are also expressed in the majority of excitatory neurons, two remaining markers (Ptger3 and Trpm2) were completely absent in scRNA-seq data analyzed with the exonic reference (Extended Fig. 3a). Reanalyzing the data with the optimized transcriptomic reference uncovered an additional 3 neuron types revealing a total of 16 neuron classes in this structure (Extended Fig. 3b). Importantly, the optimized reference also revealed one of the missing warmth activated markers – Ptger3 – that was previously missing due to intronically mapped sequencing reads (Extended Fig. 3b,c). The recovery of this gene revealed that Glut6^MnPO^ is likely the heat activated neuron type as it is labeled by all three warmth markers (Ptger3, Bdnf, Pacap). Furthermore, this analysis also revealed that three neuron classes Glut1^MnPO^, Glut2^MnPO^, and Glut3^MnPO^ are likely the thirst state encoding neurons as they are labeled by all thirst markers as well as Etv1 that was recently shown to exclusively label thirst activated neuron classes in related lamina terminalis nuclei^11^. Finally, the markers for cold and ingestion activated neurons appear to exclusively label a single cell type each (Glut4^MnPO^ for cold, Gaba1^MnPO^ for ingestion) that collectively suggests a clear mapping of previously uncovered physiological functions to the underlying neuron classes (Extended Fig. 3d,e).

Similarly, we evaluated the potential of revealing previously inaccessible cell-types and states in human peripheral blood mononuclear cell (PBMC) data through the application of optimized human transcriptomic reference. Although major cell classes in human PBMC datasets are easily detected by regular scRNA-seq analysis with exonic transcriptomic reference, this analysis fails to detect many known cell types including subclasses of CD4 and CD8 T-lymphocytes (Fig. 5f,g) ^30–32^. Moreover, the canonical markers for many known cell classes^31^ such as natural killer cells (NK), regulatory T cells (T-regs), T helper 2 cells (Th2), mucosal activated invariant T cells (MAIT) and others appear to be missing with PBMC scRNA-seq that relies on an exonic transcriptomic reference (Fig. 5f, g). By incorporating the discarded read information with our optimized human transcriptome, we were able to uncover several known cell classes within our PBMC dataset including Th17 and Th2 T-helper cells, T-regs, MAIT cells and CD8 T terminal effector cell clusters with their canonical markers as well as uncover known markers for other cell classes that are missing with exonic reference data (Fig. 5g). These data demonstrate, that incorporating intronic, 3’ intergenic and multigene mapping read data can robustly reveal biologically relevant cell types that would otherwise be obscured by poor cellular profiling resolution. In summary, optimizing transcriptomic references for 3’ scRNA-seq analysis experiments by incorporating discarded sequencing read information dramatically improves the resolution as well as biological insight from resulting analyses.

## Discussion

Previously, several approaches have been adopted to address the issue of missing genes in 3’ single cell RNA-sequencing datasets. One such approach entails using imputation to infer missing gene expression data^33^. These approaches however require deeply sequenced full-transcript scRNA-seq datasets as input, which is seldom easily available. Furthermore, imputed gene expression would have to be experimentally validated which is prohibitive from the resource vantage point. Another approach has been the usage of various pre-mRNA references, especially in case single nuclei are profiled, to capture reads mapping to unspliced pre-mRNAs^34^. We have incorporated a hybrid pre-mRNA reference strategy for registering intornic reads into our reference optimization step highlighting that the specific method for generating a pre-mRNA reference plays a major role in how many genes and reads are detected as a consequence (Extended Fig. 2). Furthermore, recent work has demonstrated that the majority of intronic reads stem from aberrant priming of transcripts from intronic poly-A tracts emphasizing the importance of not discarding these data^16,17^. Finally, a few studies have resorted to manually fixing individual loci providing a local fix to a global problem^8,11^. Here, we have undertaken a systemic effort to provide genome wide optimized transcriptomic references for mouse and human reference transcriptomes and outlined a general strategy to easily achieve the same for any genome of interest.

A surprising upshot of this study is that suboptimal use of transcriptomic references in scRNA-seq analysis impacts detection of thousands of genes for any given experiment. Our findings underscore the need to optimize genome annotations to maximize biological insight gained with this popular method. Furthermore, we showed that most exhaustive genome annotations that include gene models detailing infrequent transcripts can often be detrimental for the visibility of genes in scRNA-seq analysis. We find significantly higher number of overlapping genes in the mammalian genomes than previous estimates^35^ as genome annotations in the recent years have become more complete incorporating rarer transcript isoforms. These rare isoforms often include readthrough and premature start transcripts that severely impact read mapping in scRNA-seq analysis. One useful strategy to determine which transcript isoforms to exclude is to compare gene models between Ensembl^36^ and the somewhat more conservative Refseq^37^ genome annotations to see the level of support for rare isoforms. Furthermore, we have provided an automated classification approach to rapidly identify premature and readthrough transcripts to cut down time required for genome annotation curation. This is particularly essential as genome annotations are continuously updated.

An important consideration is to recognize that genome annotation optimization is partially dependent on the biological sample that is used to inform this process. Although resolving gene overlaps is largely independent of that, incorporating unannotated 3’ UTRs heavily relies on the input scRNA-seq dataset to identify genes with extensive intergenic read mapping proximal to their 3’ end. This is particularly important in light of recent findings that alternative splicing can significantly vary across tissues and cell types^38^ and thereby read mapping to unannotated 3’ UTRs can also vary in a cell-type specific manner^39^. Moreover, to our surprise we found significant intergenic read mapping close to 3’ ends of thousands of genes in the mouse and human genomes. While we were able to recover this sequencing information convincingly for less than a thousand genes per species, our findings point to the need for a concerted effort to update our gene models for 3’UTRs. In summary, our findings stress the importance of optimizing transcriptomic references for single-cell RNA-seq analysis, point to the need to improve genome annotations with respect to 3’ UTR models, and highlight the potential of uncovering new biology by analyzing both new and previously deposited 3’ single-cell RNA-seq datasets with optimized transcriptomic references.

## Methods

### Mice

All animal care and experimental procedures were executed in accordance with the US NIH guidance for the care and use of laboratory animals and approved by the California Institute of Technology Animal Care and Use Committee (protocol no. 1694-14). Mice were obtained from Jackson Laboratory and allowed to acclimate in the animal facility for a week. Mice were on a 12-h light–dark cycle and were provided food and water ad libitum. 6 male C57BL/6J mice at 8 weeks of age were used for microdissection of the Median Preoptic Nucleus tissue.

### Human samples

All studies were performed on human peripheral blood mononuclear cells obtained from Hemacare. The California Institute of Technology Institutional Review board (IRB) has determined that this work is exempt from the requirement for IRB review and approval (Reference #17-0727), and informed consent was not required.

### Single-cell RNA-sequencing

Single-cell RNA-sequencing library was prepared from mouse median preoptic nucleus tissue as previously described^11^. Briefly, mice were anaesthetized with isoflurane in an isolated plexiglass chamber. Brains were rapidly extracted and dropped into ice-cold carbogenated (95% O2 and 5% CO2) NMDG-HEPES-ACSF (93 mM NMDG, 2.5 mM KCl, 1.2 mM NaH2PO4, 30 mM NaHCO3, 20 mM HEPES, 25 mM glucose, 10 mM MgSO4, 1 mM CaCl2, 1 mM kynurenic-acid Na salt, 5 mM Na-ascorbate, 2 mM thiourea and 3 mM Na-pyruvate, pH adjusted to 7.4, osmolarity ranging 300–310 mOsm). Brain sections (2 mm) containing MnPO were cut with a razor blade on a stainless steel brain matrix (51392, Stoelting) and transferred to a dissection dish on ice containing NMDG-HEPES-ACSF. MnPO was microdissected with a microsurgical stab knife (72-1501, Surgical Specialties). Microdissected tissue was aggregated from 6 animals in a collection tube on ice containing NMDG-HEPES-ACSF. For enzymatic digestion of tissue, NMDG-HEPES-ACSF was replaced by trehalose-HEPES-ACSF (92 mM NaCl, 2.5 mM KCl, 1.25 mM NaH2PO4, 30 mM NaHCO3, 20 mM HEPES, 25 mM glucose, 2 mM MgSO4, 2 mM CaCl2, 1 mM kynurenic acid Na salt and 2.5 wt/vol trehalose, pH adjusted to 7.4, osmolarity ranging 330–340 mOsm) containing papain (60 U/ml; P3125, Sigma Aldrich, re-activated with 2.5 mM cysteine and a 30-min incubation at 34 °C and supplemented with 0.5 mM EDTA). Extracted MnPO tissue was incubated at 34 °C with gentle carbogenation for 45 min. During enzymatic digestion the tissue was pipetted periodically every 10 min. At the end of enzymatic digestion, the medium was replaced with 200 μl of room temperature trehalose-HEPES-ACSF containing 3 mg/ml ovomucoid inhibitor (OI-BSA, Worthington) 25 U/ml DNase I (90083, Thermo Scientific) and tissue was gently triturated into a uniform single-cell suspension with consecutive rounds of trituration with fire-polished glass Pasteur pipettes with tip diameters of 600, 300 and 150 μm. The resulting cell suspension volume was brought up to 1 ml with trehalose-HEPES-ACSF with 3 mg/ml ovomucoid inhibitor and pipetted through a 40-μm cell strainer (352340, Falcon) into a new microcentrifuge tube. Thereafter, the single-cell suspension was centrifuged down at 300g for 5 min at 4 °C and the supernatant was replaced with fresh ice-cold trehalose-HEPES-ACSF and the cell pellet was resuspended. Cells were pelleted again and resuspended in 100 μl of ice-cold resuspension-ACSF (117 mM NaCl, 2.5 mM KCl, 1.25 mM NaH2PO4, 30 mM NaHCO3, 20 mM HEPES, 25 mM glucose, 1 mM MgSO4, 2 mM CaCl2, 1 mM kynurenic acid Na salt and 0.05% BSA, pH adjusted to 7.4, osmolarity ranging 330–340 mOsm). Cell suspension volumes estimated to retrieve ∼6,000 single-cell transcriptomes were added to the 10x Genomics RT reaction mix and loaded to the 10X Single Cell A chip (230027, 10x Genomics) per the manufacturer’s protocol. We used the Chromium Single Cell 3’ Library and Gel Bead Kit v2 (120237, 10x Genomics) and the Chromium i7 Multiplex Kit (120262) to prepare Illumina sequencing libraries.

Human peripheral blood mononuclear cell scRNA-seq libraries were prepared as prevously reported^40^. Briefly, human cryopreserved PBMCs were thawed in 37 °C RPMI-1640 medium and pelleted at 300 × *g* for 2 min. Cells were resuspended to 1e^6^ cells/mL in RPMI-1640. Cells estimated to retrieve 10 000 single-cell transcriptomes were loaded into a 10X Genomics lane using single-cell 3′ v2 reagents following manufacturer’s protocol.

### Sequencing, read-mapping and generation of digital expression data

scRNA-seq sequencing libraries were sequenced on an Illumina HiSeq 4000 sequencer. Mouse sequencing data were aligned to either the latest mouse exonic transcriptomic reference provided by 10x Genomics (mouse GRCm38 primary sequence assembly and Ensembl 98 genome annotation) or optimized transcriptomic reference based on the latter using Cell Ranger 6.1.2 (10x Genomics) or STARsolo (STAR 2.7.9a). Cell Ranger *count* pipeline was run under default parameters to generate exonic read mapping based gene-cell matrices. STARsolo readmapping pipeline was run with the following specified parameters: *--clipAdapterType CellRanger4 -- outFilterScoreMin 30 --soloCBmatchWLtype 1MM_multi_Nbase_pseudocounts --soloUMIfiltering MultiGeneUMI_CR --soloUMIdedup 1MM_CR --outSAMtype BAM SortedByCoordinate -- readFilesCommand zcat --soloType CB_UMI_Simple --bamRemoveDuplicatesType UniqueIdenticalNotMulti --outSAMattributes NH HI AS nM GX CB UB GN sM sQ --soloFeatures GeneFull --soloCBwhitelist /<location folder>/737K-august-2016.txt.* Intronic data was incorporated by specifying *--include-introns* parameter in Cell Ranger or *--soloFeatures GeneFull* in STARsolo. Human sequencing data was aligned to the latest human exonic transcriptomic reference provided by 10x Genomics (based on human GRCh38 primary sequence assembly and Ensembl 98 genome annotation) or optimized reference based on the latter and processed as above.

## Data analysis

### Estimation of exonic, intronic and intergenic read fractions

Read idenitiy (exonic, intronic or intergenic) of confidently mapped reads was evaluated with the Genomic Alignments (1.28.0) R package. Aligned reads from the Cell Ranger output bam file from sequencing data mapped to the regular exonic reference were imported with RE, NH and AN tags including duplicate reads and excluding secondary alignments (*param = ScanBamParam(flag = scanBamFlag(isSecondaryAlignment = FALSE*). Read itentity proportions were estimated from all non-secondary uniquely mapped reads (NH=1). The number of exonic reads was estimated by extracting all reads classified as exonic (RE=“E“) and subtracting all exonic reads with registered mapping to antisense reads (AN!=NA). Removing exonic reads with antisense gene annotations was necessary since Cell Ranger 6 and earlier versions wrongly classify intergenic reads mapping antisense to known exons as exonic. Intronic read number was obtained by identifying reads classified as intronic (RE=“N“). Finally, the total number of intergenic reads was obtained by adding reads classified as intergenic (RE=“I“) to false exonic reads mapped antisense to known exons (RE=“E“ & AN!=NA).

### Determining dominant source of read mapping for genes

In order to determine which fraction of genes is detected mostly by exonically, intronically or intergenically mapped reads we estimated exonic, intronic and 3’ intergenic read counts within 10kb of known gene ends for each gene. To determine the number of exonic reads for each gene we extracted all reads annotated to a specific gene from the raw gene-cell matrix from scRNA-seq dataset mapped to the exonic reference with Cell Ranger 6.1.2. Intronic reads mapping to each gene were estimated by subtracting the exonic read estimates from gene expression estimates obtained from mapping scRNA-seq data to the traditional pre-mRNA reference where full transcripts are defined as exons. Finally, 3’ intergenic read estimates within 10kb of known gene ends were obtained by extracting all intergenic reads (RE=I as well as RE=E & AS!= NA as Cell Ranger misclassifies reads mapping antisense to exons as exonic, whereas they are mostly intergenic reads) from the Cell Ranger genome aligned sequenging data .bam file with GenomicAlignments R package. Only unique intergenic reads with non-defective molecular and cellular barcodes were retained and ascribed to the closest 3’ end of a gene with bedtools. All intergenic reads further than 10kb from known gene ends were filtered out with the rest used to estimate 3’ intergenic read mapping for each gene. Dominant source of read mapping was estimated for all genes with more than 100 reads mapping per locus with exonic dominant genes with more than 50% of reads stemming from exons, intronic dominant genes with more than 50% of reads stemming from introns and intergenic dominant genes with more than 50% of reads stemming from within 10kb 3’ of known gene ends.

### Evaluation of intergenic read distribution at 3’ gene ends

3’ intergenic read data was extracted from Cell Ranger genome aligned sequenging data .bam file with GenomicAlignments R package (intergenic reads were identified by read tag RE=I). Extracted reads were ascribed to the closest 3’ end of a gene with estimated distance from respective gene end with bedtools and resulting intergenic read distribution from gene ends was plotted.

### Evaluation of intronic read incorporation with different pre-mRNA references

In order to evaluate different strategies for incorporating intronic reads in scRNA-seq analysis, we evaluated gene and sequencing read detection with three independent approaches. (a) in “Cell Ranger pre-mRNA reference” we compiled a new genome annotation file by transferring all “transcript” feature entries from the original genome annotation (GENCODE vM23/Ensembl 98 for mouse data and GENCODEv32/Ensembl 98 for human data) to a new gene transfer format (gtf) file and renaming the feature field for all entries as “exon” effectively redefining all full length transcripts as exons and thereby enabling registration of intronic reads. We compiled a new Cell Ranger transcriptomic reference with Cell Ranger 6.1.2 using the *cellranger mkref* pipeline with default parameters and applied the newly assembled transcriptomic reference in sequencing read mapping with *cellranger count* pipeline again with standard parameters (see https://github.com/PoolLab/Generecovery for details); (b) for “STARsolo GeneFull mode” we generated the STAR reference transcriptome using STAR 2.7.9a in genomeGenerate run mode with mouse or human genome sequence (mouse GRCm38 genome build or human GRCh38 genome build, respectively) and corresponding genome annotation using default parameters. Sequencing read data was then mapped to the generated STAR reference transcrtiptome using STARsolo with the following parameters: *STAR --genomeDir <location of transcriptomic reference> --readFilesIn <location of sequencing fastq files> --clipAdapterType CellRanger4 -- outFilterScoreMin 30 --soloCBmatchWLtype 1MM_multi_Nbase_pseudocounts --soloUMIfiltering MultiGeneUMI_CR --soloUMIdedup 1MM_CR --outSAMtype BAM SortedByCoordinate -- readFilesCommand zcat --soloType CB_UMI_Simple --bamRemoveDuplicatesType UniqueIdenticalNotMulti --outSAMattributes NH HI AS nM GX GN sM sQ CB UB --soloFeatures GeneFull --soloCBwhitelist <location of barcode whitelist>/737K-august-2016.txt*. Note that specifying “FullGene” mode results in intronically mapped reads being incorporated in the gene-cell matrix. Finally,in “Cell Ranger --include-introns mode” (c), we used the previously generated Cell Ranger transcriptomic reference to run the *cellranger count* pipeline with *--include-introns* parameter specified resulting in inclusion of intronic reads in the resulting gene-cell matrix.

In order to compare gene detection fidelity between the above approaches, we imported raw unfiltered gene-cell matrices generated by Cell Ranger or STAR into Seurat 4.1.0 and identified all expressed genes from raw gene expression data. Unique and shared gene sets were identified by intersection of individual gene sets in R. We compared the read detection efficiency for the three strategies by extracting individual sequencing reads from Cell Ranger and STAR generated BAM files with GenomicAlignments 1.28.0 package in R. For Cell Ranger generated BAM files we extracted non-duplicate reads with *readGAlignments(bamfile, index=indexfile, param = ScanBamParam(flag = scanBamFlag(isDuplicate = FALSE, isSecondaryAlignment = FALSE), tag = c(“GN”, “RE”, “xf”, “CB”, “UB”), what = “flag”)).* We identified all reads contributing to the gene-cell matrix in Cell Ranger BAM files by pulling out reads where *xf==25* and extracted their unique molecular and cellular barcodes by combining their CB and UB fields. For STAR BAM files we imported sequencing read data with readGAlignments(bamfile, index=indexfile, param = ScanBamParam(flag = scanBamFlag(isDuplicate = FALSE, isSecondaryAlignment = FALSE), tag = c(“GX”, “GN”, “CB”, “UB”), what = “flag”)). Reads contributing to gene-cell matrices in STAR BAM files were identified by pulling out reads where *GX!=NA*, and length of cell and UMI barcodes were and 10 characters, respectively. Unique and shared detected reads was evaluated by intersecting the combined CB/UB indeces of reads between different methods.

### Assembling optimized transcriptomic references

We took a three pronged approach to generating a scRNA-seq optimized transcriptomic reference which involved the following steps: a) resolving gene overlap derived read loss; b) recovering intergenic reads from 3’ unannotated exons; and c) recovering intronic reads. Each of these steps generated input data for a custom R script for automated assembly of the scRNA-seq optimized genome annotation.

#### Resolving gene overlap derived read loss

We generated a custom code (available at https://github.com/PoolLab/Generecovery) to identify same-strand overlapping genes in a given input genome annotation and triage genes for manual curation to resolve the overlap derived sequencing read loss. Overlapping genes were identified by evaluating chromosomal location (*seqname, start* and *end* coordinates as well *strand* in the gene transfer format file for all “gene” features) based on which we generated an output file summarizing the overlapping gene list specifying the number of overlapping genes and their identity. Gene overlaps can be resolved by one of several strategies including (i) leaving overlapping gene annotations unchanged if their exons don’t directly overlap, (ii) deleting offending readthrough transcripts from upstream genes, (iii) deleting offending premature gene transcripts from downstream genes, (iv) deleting pseudogenes and non-protein-coding genes with poor support and no read mapping that obscure well established protein-coding genes or (v) for extensively overlapping genes deleting one and renaming the other to capture otherwise discarded reads. As well annotated genomes contain several thousand same-strand overlapping genes and properly resolving gene overlaps often requires manual inspection of the locus to determine best course of action, prioritization of genes for manual curation is often desirable. To this end, we generated a gene classification algorithm to prioritize genes for direct inspection. The following algorithm was used to classify genes for appropriate curation:

1. If gene overlaps with several genes:
  a. Classify for “Manual inspection” if multigene overlapping gene as well as its corresponding nested genes have overlapping exons.
  b. Classify as “keep as is” if multigene overlapping gene’s individual exons do not overlap.
    i. If nested gene does not overlap with any other gene, classify as “Keep as is”
    ii. If nested gene overlaps with more than one gene, classified for “Manual inspection”
2. If gene overlaps with only one other gene, test whether gene is non-protein-coding/pseudogene (“Gm” and “…Rik” gene models in mice; “AC…” and “AL…” gene models in humans)
  a. If both overlapping genes are non-protein-coding/pseudogenes, classify for “Manual inspection”
  b. If only one gene in the overlapping gene pair is non-protein-coding/pseudogene, test if genes have overlapping exons:
    i. In case no overlapping exons, classify both genes as “Keep as is”
    ii. In case exons overlap, mark non-protein-coding/pseudogene for deletion (“Delete”).
  c. If both genes are well supported genes:
    i. If their exons don’t overlap, mark both genes as “Keep as is”
    ii. If their exons do overlap, determine the number of opposing gene’s exonic overlap for each exon of each gene and find the exon with most overlaps for both upstream and downstream gene to determine appropriate course of action:
      1. If downstream gene’s exon has more overlaps than its upstream counterpart, classify downstream gene as “Premature transcript deletion” and upstream gene as “Keep as is”
      2. If upstream gene’s exon has more overlaps than its downstream counterpart, classify upstream gene as “Readthrough transcript deletion” and downsream gene as “Keep as is”.
      3. Otherwise classify both for “Manual inspection”

The resulting recommendations were used in the manual curation step, where all genes that were not classified in the “Keep as is” category were directly scrutinized in the Ensembl genome browser (ensemble.org, with GRCm38 and GRCh38 genome builds for mouse and human data, respectively) and/or cross-referenced to the respective Refseq genome annotation within the Integrated Genome Browser (IGV 2.11.9). For genes with multiple transcripts, readthrough and premature transcripts were marked for deletion in a separate column in the overlapping gene list file if the remaining transcripts enabled registration of reads from core exons of both genes. In the few cases where the terminal exons of genes were completely overlapping we deleted one of the genes and marked the other for renaming to avoid discarding expression data from both genes. For genes marked for deletion, we cross-referenced the gene structure to Refseq annotation and confirmed the deletion if the gene model was not supported by both Ensembl and Refseq genome annotations and the gene overlap could not be resolved by deleting a specific transcript. For candidate genes for deletion we also confirmed that the gene models in question would not have any reads mapping to their loci with the mouse brain and human PBMC datasets. The resulting curated ‘overlapping gene list’ file with final action recommendations and transcript identities for deletion was saved and used as input in the automated generation of optimized genome annotation.

#### Recovering intergenic reads from 3’ unannotated exons

Gene detection in 3’ single-cell RNA-sequencing depends on registering sequencing reads predominantly at the 3’ end of transcripts. Therefore, inaccurate annotation of 3’ exons and UTRs causes misclassification of exonic sequencing reads as intergenic and thereby compromised detection of gene expression. In order to identify candidate genes with poorly annotated 3’ regions for manual correction of gene boundaries we identified and rank-ordered genes with high numbers of intergenic sequencing reads mapping within 10kb of known gene ends. This candidate gene list was stored in a ‘gene extension candidates’ output file and subjected to case-by-case scrutiny for evidence for unannotated 3’ exons with Integrated Genomics Viewer (IGV 2.11.9). New estimates for 3’ gene boundaries were recorded in the ‘gene extension candidates’ file for genes displaying evidence of unannotated exons. Evidence for the latter included sequencing read splicing between known and putative unannotated exons, continuous sequencing read mapping extending 3’ of known gene end or independent expression validation with an independent method (e.g. Human Protein Atlas and Allen In Situ Mouse Brain Atlas datasets).

To achieve this, we mapped mouse MnPO and human PBMC scRNA-seq data to their respective reference transcriptomes with Cell Ranger and imported all intergenic reads from aligned read data (bam file) with GenomicRanges R package using *readGAlignments(bamfile, index=indexfile, param = ScanBamParam(flag = scanBamFlag(isDuplicate = FALSE, isSecondaryAlignment = FALSE), tag = c(“GN”, “RE”), what = “flag”, tagFilter = list(“RE”=“I”)))*. Duplicate and defective intergenic reads were removed by retaining reads with unique barcodes with 10x defined correct length of cellular and molecular barcodes. We generated a bed file of the intergenic reads using the GenomicRanges and rtracklayer R packages. Next we generated the reference bed file with gene boundaries by extracting all *gene* feature entries form the genome annotation (gtf file) and reformatted to match the bed file requirements. We used bedtools (2.30.0) to identify the closest 3’ gene end to all intergenic reads with *bedtools closest -a intergenic_reads.bed -b gene_ranges.bed -s -D a -fu > <results>.txt*. We used these data to estimate the total number of all intergenic reads within 10kb of each gene, rank ordered genes from highest to lowest number of intergenic reads proximal to 3’ end and saved the resulting data to ‘gene extension candidates’ file. We proceeded to scrutinize the resulting gene list and evaluated evidence for unannotated 3’ exons by visualizing read mapping and splicing data in the aligned sequencing data (bam file) with Integrated Genomics Viewer following the criteria above. For genes where these criteria were satisfied we recorded the new extended gene boundary in the ‘gene extension candidates’ file. Furthermore, we extracted genomic coordinates for 3’ gene ends from the Refseq annotation for all mouse and human genes and updated 3’ gene ends to reflect Refseq coordinates if Refseq genome annotation placed the gene end 3’ of the Ensembl annotation. The resulting ‘gene extension candidates’ file was used as input in the automated generation of optimized genome annotation.

#### Recovering intronic reads

All currently available strategies for incorporating intronic reads come with their own limitations and can differentially detect several hundred genes. The traditional approach for generating a pre-mRNA reference entails redefining all transcripts as exons. Despite robustly increasing overall registered reads in the resulting gene-cell matrix, this approach significantly exacerbates gene overlap derived read loss and fails to detect many spliced intronic reads due to eliminating known exon boundaries. Later versions of sequencing read aligners allow specific intronic mapping modes such as specifying *--include-introns* attribute in Cell Ranger. However, although this latter method circumvents issues with gene overlap derived read loss common for traditional pre-mRNA references and it outperforms the traditional pre-mRNA reference approach by detected read number, surprisingly this solution fails to detect many intronic sequencing reads that are commonly detected by the latter. Therefore we adopted a hybrid approach to capturing intronic reads by generating a reference transcriptome that incorporates both normal exonic gene structures as well as full transcript length exons that enabled us to effeciently capture all intronic reads under Cell Ranger *--include-introns* mode. To this end we generated the traditional pre-mRNA reference genome annotation by transferring all *transcript* feature entries from the genome annotation gtf file and redefinined their feature value as *exon* saving it as a new ‘traditional pre-mRNA annotation’ file that we used as input in the final generation of the scRNA-seq optimized genome annotation assembly.

#### Automated generation of the scRNA-seq optimized genome annotation

The following input data was used to generate the scRNA-seq optimized genome annotation: (i) regular genome annotation (.gtf file containing Ensembl v98 genome annotations from 10x Genomics for the respective genome build), (ii) ‘traditional pre-mRNA annotation’ (.gtf file), (iii) ‘overlapping gene list’ file specifying transcripts and genes for deletion and (iv) ‘gene extension candidates’ file specifying updated 3’ gene boundaries for genes with unannotated 3’ UTRs. We wrote an R script for assembling the final scRNA-seq optimized genome reference (available at https://github.com/PoolLab/Generecovery). Briefly, we imported the regular genome annotation with rtracklayer R package, removed genes and transcripts marked for deletion in the ‘overlapping gene list’ file, adjusted 3’ gene boundaries for genes specified in the ‘gene extension candidates’ file, imported the ‘traditional premRNA annotation’ data and added full transcript length exonic annotations to all genes excluding genes in the overlapping gene list, renamed defined genes and exported the resulting genome annotation to a new <scRNA-seq optimized genome annotation.gtf> file. The resulting genome annotation thus eliminates the structural problems of existing genome annotations that unnecessarily discard sequencing read data that reflect true endogenous gene expression.

#### Assembly and use of the scRNA-seq optimized transcriptomic reference

The resulting <scRNA-seq optimized genome annotation.gtf> was used to compile the optimized transcriptomic reference with Cell Ranger using the *cellranger mkref* pipeline. The resulting optimized transcriptomic reference was used to align sequencing data with the *cellranger count* pipeline with *--inculde-introns* attribute specified.

### Data analysis and visualization software

scRNA-seq data were processed in R 4.1.2 using the following analysis and data visualization packages: RStudio 1.4.1717, Seurat v.4.1.0., ggplot2 3.3.5, GenomicAlignments 1.28.0, GenomicRanges 1.44.0, IRanges 2.26.0, rtracklayer 1.52.1, Matrix 1.3-4, readxl 1.3.1 and stringr 1.4.0. Gene overlaps in genome annotations and overlapping gene classification code was generated by Python3 and R. We used bedtools 2.30.0 to identify intergenic gene reads proximal to known 3’ gene ends.

Reference transcriptomes for sequencing read mapping were generated by either Cell Ranger 6.1.2 or STARsolo (STAR 2.7.9) using *mkref* and *genomeGenerate* pipelines, respectively. Sequencing data were mapped to dedicated reference transcripomes with either Cell Ranger 6.1.2 *count* pipeline or STARsolo as described above.

Sequencing read mapping data in Cell Ranger aligned BAM files was visualized and evaluated with Integrated Genomics Viewer 2.11.9. Read mapping data with respect to specific gene exons and introns was plotted in R with Sushi (1.30.0) package. Transcript and gene structures were plotted with Gviz 1.36.2 R package. We used Graphpad Prism 8 to plot summary data.

## Supporting information

Supplemental Table 1 - Mouse Genome Annotation Updates

Supplemental Table 2 - Human Genome Annotation Updates

## Data Availability

Raw and processed scRNA-seq data are available at the NCBI Gene Expression Omnibus (GEO accession no. GSE198528). Latest versions of the mouse and human reference transcriptomes and genome annotations are available for download at www.thepoollab.org/resources.

## Code Availability

Code to reproduce data analysis in this manuscript and generate scRNA-seq optimized genomic annotations is available at https://github.com/PoolLab/Generecovery.

## Acknowledgements

We thank Lior S. Pachter and members of the M.T. lab for helpful discussion and comments. We thank the Single-Cell Profiling Center (SPEC) in the Beckman Institute at Caltech for technical assistance with scRNA-seq. A.H.P. is supported by Eugene McDermott Scholar funds and by Startup funds from Peter O’Donnell Jr. Brain Institute at UT Southwestern. Y.O. is supported by Startup funds from the President and Provost of the California Institute of Technology and the Biology and Biological Engineering Division of California Institute of Technology, Searle Scholars Program, the Mallinckrodt Foundation, the McKnight Foundation, the Klingenstein-Simons Foundation, the New York Stem Cell Foundation and the NIH (R56MH113030 and R01NS109997).

## Author Contributions

A.H.P. conceived and designed the project. A.H.P. and H.P. devised and performed data analysis. A.H.P. and S.C. generated the MnPO scRNA-seq dataset. S.C. and M.T. generated the human PBMC scRNA-seq dataset. S.C., M.T. and Y.O. provided conceptual advice on data analysis. All authors contributed to the manuscript as drafted by A.H.P. and H.P. A.H.P. and Y.O. supervised the overall project.

## Supplemental figures

**Extended data Figure 1:**
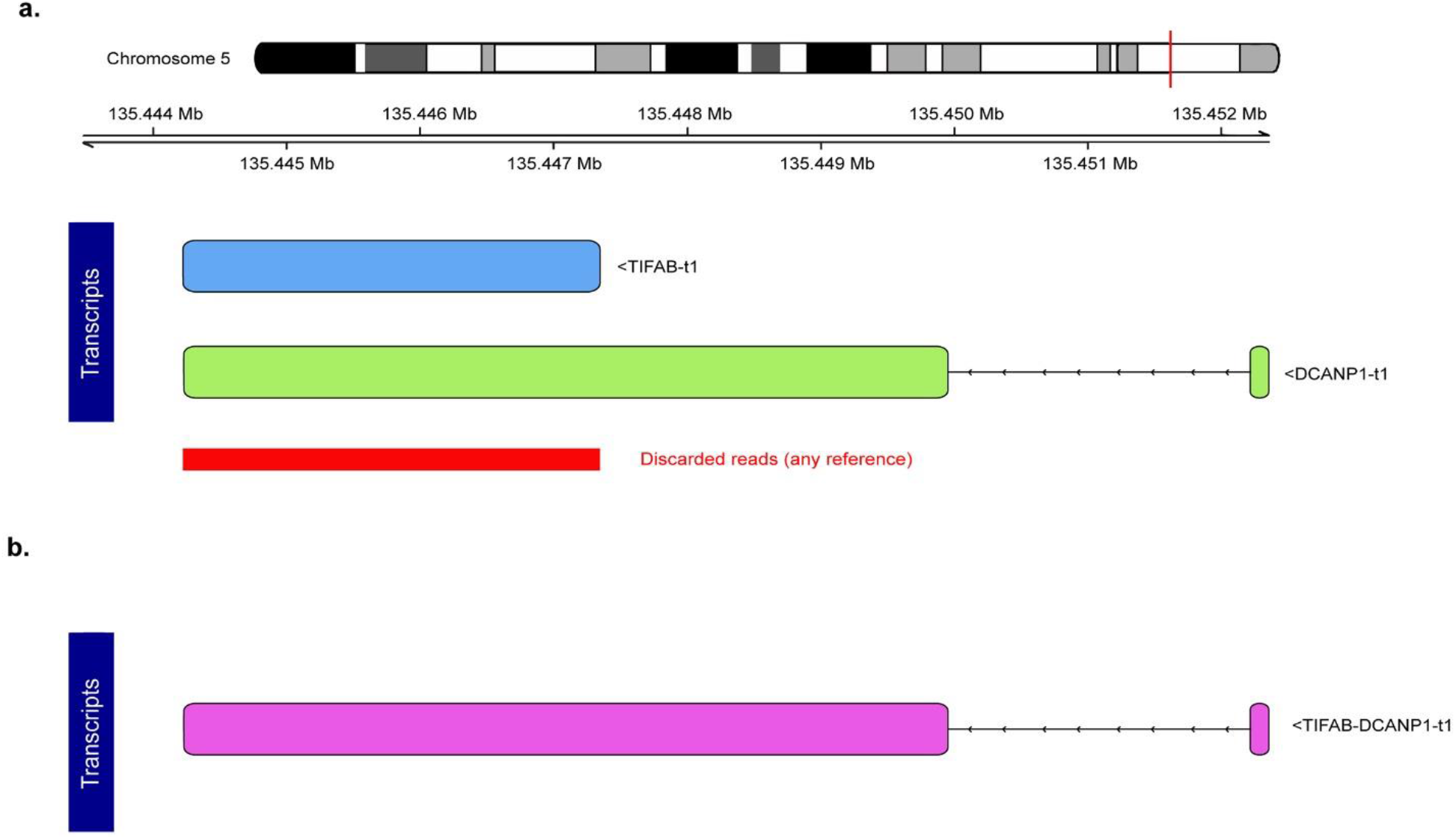
Genes with shared terminal exon sequences are obscured from scRNA-seq analysis. **a.** The terminal exons of human genes TIFAB and DCANP1 overlap resulting in all sequencing reads mapping to the overlapping area being discarded due to “multigene mapping” classification. Thereby, 3’ scRNA-seq is mostly blind to the expression of these genes. **b.** Expression information for TIFAB and DCANP1 genes can be recovered by removing one of the genes and renaming the remaining gene’s transcript recovering discarded expression data.

**Extended data Figure 2:**
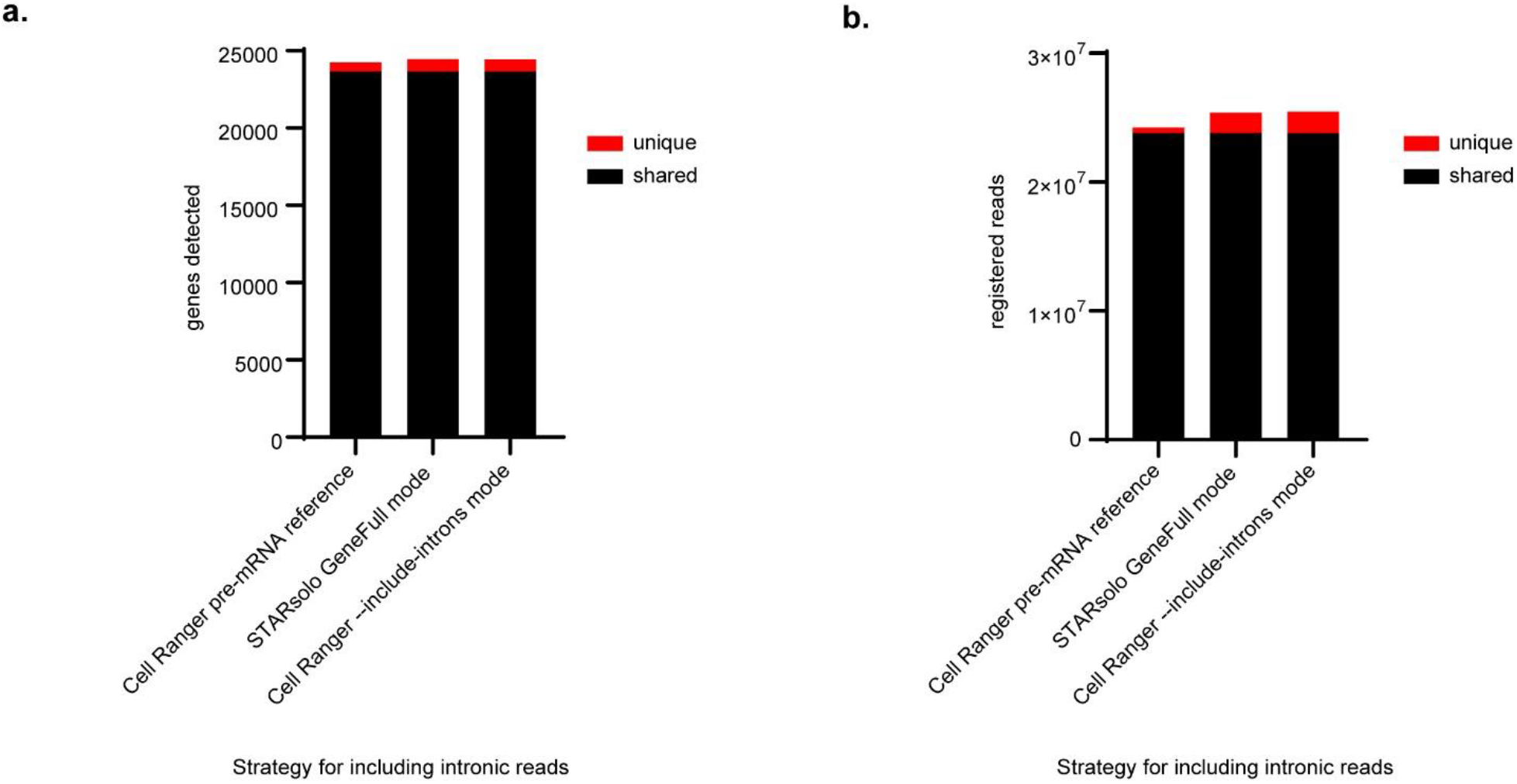
Different strategies for incorporating intronic reads into scRNA-seq analysis varie by detection of several hundred genes and up to 6.5% of sequencing reads. **a.** Comparison of detected genes by distinct methods for incorporating intronic sequencing reads into scRNA-seq analysis with mouse MnPO brain nucleus dataset. “Cell Ranger pre-mRNA reference” data was generated with Cell Ranger software in regular mapping mode using a genome annotation where all transcripts were defined as exons thus leading to incorporation of previously intronically mapping reads. “STARsolo GeneFull mode” data was generated by the STARsolo software by specifying the “GeneFull” attribute integrating intronically mapped reads into the assembled gene-cell matrix. “Cell Ranger --include-introns mode” was generated with the Cell Ranger software with “--include-introns” parameter specified leading to integration of intronic reads to assembly of the gene-cell matrix. Black – genes detected by all three methods for incorporating intronic reads; red – genes that are either unique or shared by only one other mehod for detecting intronic reads due to gene overlaps or problems with incorporating a subset of intronic reads with one or more of the methods. **b.** Comparison of detected sequencing reads incorporated into the gene-cell matrix by distinct methods registering intronic sequencing reads into scRNA-seq analysis. Black - reads detected by all three methods; red - reads that are either uniquely detected by a given method or shared with only one other strategy.

**Extended data figure 3:**
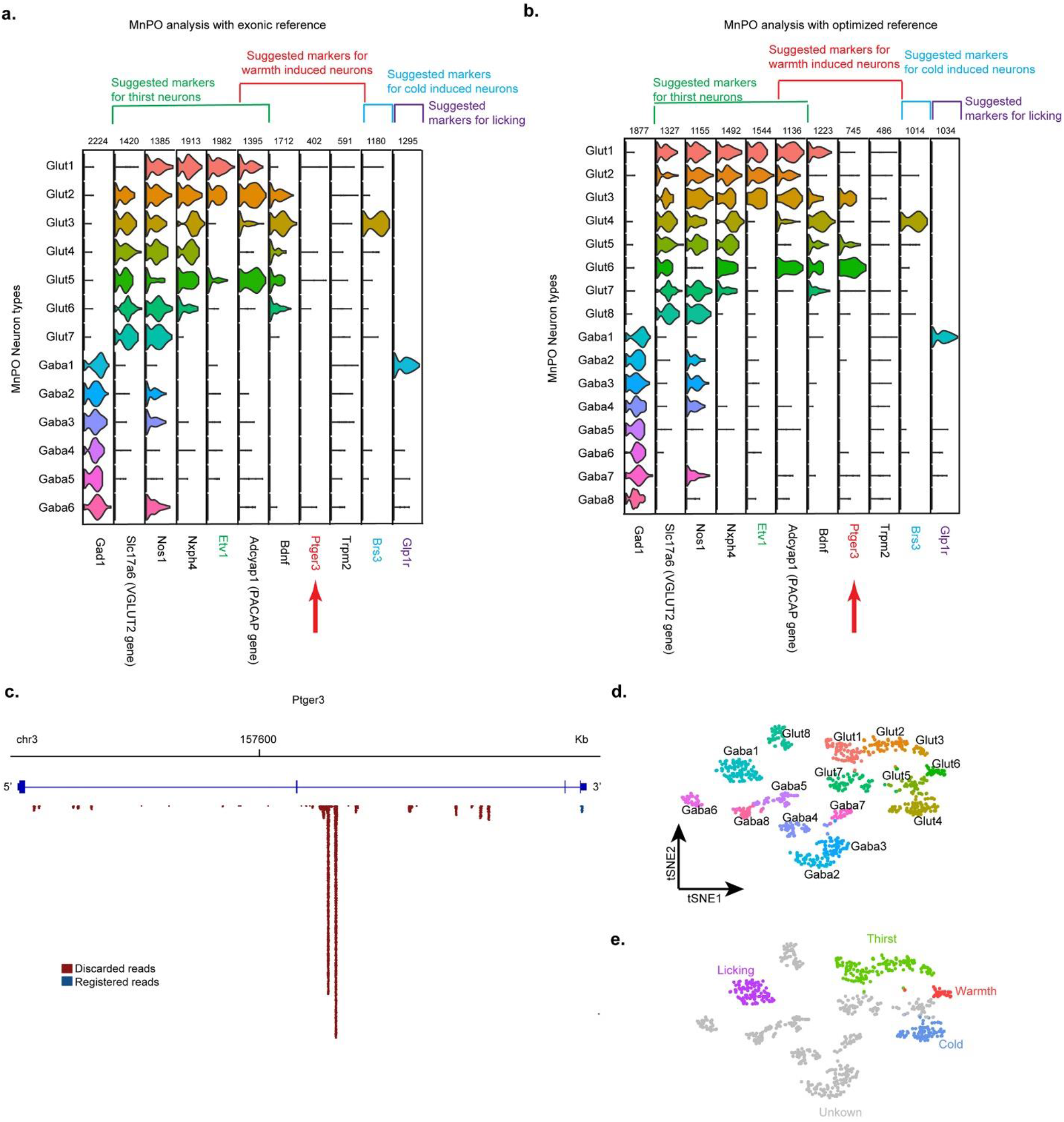
Elucidation of neuron types, cell-type-specific markers and physiological functions of cells in the mouse Median Preoptic Nucleus (MnPO) with regular exonic and optimized transcriptomic reference based scRNA-seq analyses. **a.** Violin plot of the expression of previously implicated genetic markers labeling thirst, warmth, cold or licking activated neurons in the MnPO as analyzed by a regular exonic reference based analysis pipeline. Red arrow – absence of Ptger3 – a marker for warmth activated neurons in the MnPO – in the scRNA-seq dataset. Expression is shown on a log-normalized scale with maximum counts per million (max CPM). **b.** Violin plot of the same marker expression in the MnPO as analyzed with the optimized transcriptomic reference pipeline. Red arrow – detected warmth activated neuronal marker Ptger3 expression. **c.** Mapping of sequencing reads to the Ptger3 locus with the majority being discarded from downstream analysis with an exonic reference due to their intronic mapping. **d.**Nomenclature of neuron types in the MnPO as identified by scRNA-seq with the optimized transcriptomic reference (data same as in Figure 5b). **e.** Suggested function to cell-type mapping in the MnPO neurons based on overlap of previously identified genetic markers in the new neuronal nomenclature.

